# CD4 downregulation precedes Env expression and protects HIV-1-infected cells from ADCC mediated by non-neutralizing antibodies

**DOI:** 10.1101/2024.05.01.592003

**Authors:** Jonathan Richard, Gérémy Sannier, Li Zhu, Jérémie Prévost, Lorie Marchitto, Mehdi Benlarbi, Guillaume Beaudoin-Bussières, Hongil Kim, Yaping Sun, Debashree Chatterjee, Halima Medjahed, Catherine Bourassa, Gloria-Gabrielle Delgado, Mathieu Dubé, Frank Kirchhoff, Beatrice H. Hahn, Priti Kumar, Daniel E. Kaufmann, Andrés Finzi

## Abstract

HIV-1 envelope glycoprotein (Env) conformation substantially impacts antibody-dependent cellular cytotoxicity (ADCC). Envs from primary HIV-1 isolates adopt a prefusion “closed” conformation, which is targeted by broadly-neutralizing antibodies (bnAbs). CD4 binding drives Env into more “open” conformations, which are recognized by non-neutralizing Abs (nnAbs). To better understand Env-Ab and Env-CD4 interaction in CD4+ T cells infected with HIV-1, we simultaneously measured antibody binding and HIV-1 mRNA expression using multiparametric flow cytometry and RNA-flow fluorescent *in situ* hybridization (FISH) techniques. We observed that *env* mRNA is almost exclusively expressed by HIV-1 productively-infected cells that already downmodulated CD4. This suggest that CD4 downmodulation precedes *env* mRNA expression. Consequently, productively-infected cells express “closed” Envs on their surface, which renders them resistant to nnAbs. Cells recognized by nnAbs were all *env* mRNA negative, indicating Ab binding through shed gp120 or virions attached to their surface. Consistent with these findings, treatment of HIV-1 infected humanized mice with the ADCC mediating nnAb A32 failed to lower viral replication or reduce the size of the viral reservoir. These findings confirm the resistance of productively-infected CD4+ T cells to nnAbs-mediated ADCC and question the rationale of immunotherapy approaches using this strategy.

**Graphical abstract:** 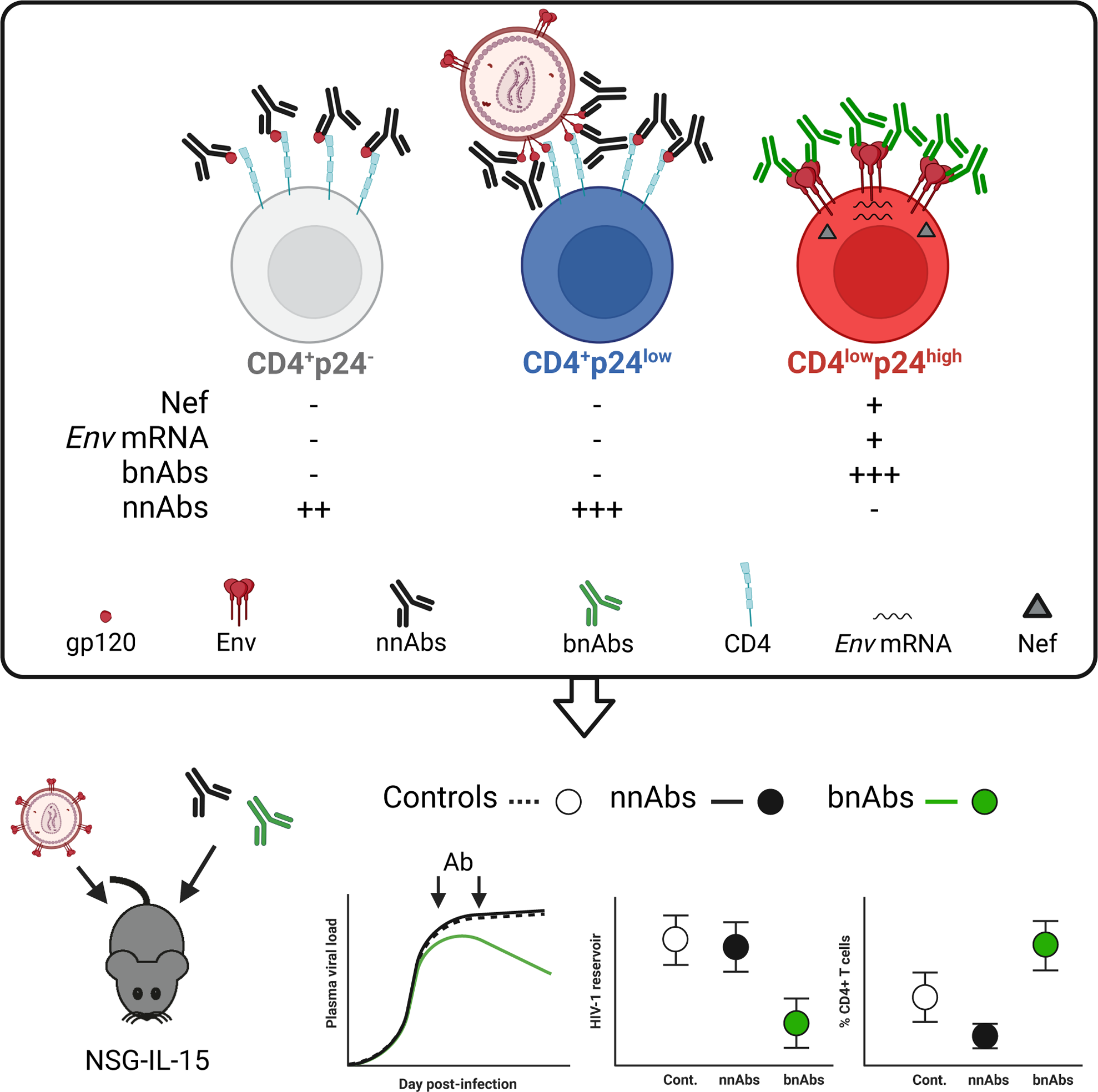

## INTRODUCTION

Highly active antiretroviral therapy (ART) efficiently suppresses HIV-1 replication and significantly increase the life expectancy of people living with HIV-1 (PLWH) (1, 2). However, it has become evident that even lifelong ART cannot eradicate the virus. Viral reactivation and rebound can occur upon treatment interruption due to the presence of a latent reservoir, persisting mainly in long-lived memory CD4+ T cells (3–6). New approaches aimed at eradicating or functionally curing HIV-1 infection by targeting and eliminating productively or latently-infected cells are needed. Monoclonal antibodies (mAbs) are attractive therapeutics for HIV cure strategies, since they target virus specific antigens and have the potential to harness host immune responses such as antibody-dependent cellular cytotoxicity (ADCC). The HIV-1 envelope glycoprotein (Env) is the only viral antigen that is present on the surface of HIV-1-infected cells, thus representing the main target for immunotherapy-based strategies (7). In recent year, several clinical trials explored broadly-neutralizing antibodies (bnAbs) as therapeutic agents to reduce the HIV-1 reservoir via Fc-mediated effector functions (8). Monotherapy or combination of bnAbs targeting multiple regions of the Env trimer including the V3 glycan supersite (10-1074, PGT121), the V2 apex (PGDM 1400, CAP256-VRC26), or the CD4 binding site (CD4bs) (3BNC117, N6, VRC01, VRC07-523) are currently under investigation. Administration of bnAbs to humans has been found to be safe and effective in lowering viremia and maintaining viral suppression for varying periods of time after treatment interruption (9–15). However, recent data suggest that bnAbs may not be as broad and/or effective as predicted, in part because of circulating viruses with pre-existing resistance to the administered bnAbs (8). So-called “non-neutralizing” Abs (nnAbs) have been evaluated as a potential alternative, because they target highly conserved Env epitopes that are usually occluded in the closed trimer, including the coreceptor-binding site (CoRBS) (16–18), the inner domain of gp120 (19–21) or the gp41 immunodominant domain (gp41ID) (22). These nnAbs are naturally elicited during HIV-1 infection and can mediate potent Fc effector functions (22–28).

HIV-1 Env is a conformationally flexible molecule that transitions from an unliganded “closed” State 1 conformation to an “open” CD4-bound State 3 conformation (29–31) upon CD4 interaction. Envs from most primary HIV-1 isolates adopt a “closed” conformation that is efficiently targeted by bnAbs, but is resistant to nnAbs (29, 32–36), which target epitopes that are occluded within the unliganded “closed” trimer. Env interaction with CD4 or small CD4-mimetic compounds are known to induce more “open” conformations, thus sensitizing infected cells to ADCC mediated by CD4-induced (CD4i) nnAbs (23, 24, 26, 27, 33, 34, 36–42). To avoid exposing these nnAb epitopes, the Env trimer is stabilized by multiple intermolecular interactions, including between the V1, V2 and V3 variable loops as well as the gp120 β20–β21 element, which maintain a “closed” conformation (31, 43, 44). In addition, HIV-1 expresses the accessory proteins Nef and Vpu, which indirectly modulate Env conformation by downregulating CD4 from the surface of infected cells, thus preventing a premature Env-CD4 interaction that would otherwise result in the exposure of CD4i Env epitopes (23, 26). Vpu-mediated counteraction of the restriction factor BST-2 (also named “tetherin”), known to tether viral particles at the surface of infected cells, also reduces cell-surface levels of Env (23, 26, 45, 46).

Despite advances in understanding Env-Ab and Env-CD4 interactions, the capacity of nnAbs to target and eliminate productively-infected cells by ADCC remain controversial. Several studies reported that cells infected with primary viruses expressing functional Vpu and Nef proteins are resistant to ADCC mediated by CD4i nnAbs, because they maintain surface expressed Env in a “closed” conformation (23, 24, 27, 32–42, 47, 48). Other studies report that CD4i nnAbs can mediate ADCC responses, but they have used infectious molecular clones (IMCs) that are defective for Nef expression (49–60). These IMCs contain reporter genes (e.g., the Renilla luciferase [LucR] gene) upstream of the *nef* gene, which significantly reduces Nef expression and thus CD4 downregulation from the surface of infected CD4+ target cells. The resulting premature Env-CD4 interaction promotes the artificial exposure of otherwise occluded CD4i epitopes (39, 61, 62).

Here, we used multiparametric flow cytometry and RNA-flow fluorescent *in situ* hybridization (FISH) techniques to characterize cell populations targeted by bNAbs and nnAbs in the context of primary CD4+ T cell infection. We show that *env* mRNA is specifically detected among cells that already downmodulated cell-surface CD4, which renders these cells refractory to recognition by nnAbs. Although some CD4+ cells are bound by nnAbs, they are all negative for *env* mRNA, indicating that they are not productively-infected and nnAbs binding is mediated through the recognition of shed gp120 and/or viral particles coated at their cell surface. The same results were obtained with *ex vivo*-expanded CD4+ T cells isolated from PLWH. Finally, the ADCC mediating nnAb A32 failed to reduce HIV-1 replication and the size of the viral reservoir in a humanized mouse model (hu-mice) that supports Fc-effector functions. These findings raise questions about curative immunotherapy-based strategies that rely solely on nnAbs, specifically those targeting CD4i epitopes.

## RESULTS

### Expression of CD4, p24 and Nef defines recognition of HIV-1 infected CD4+ T cells by nnAbs and bnAbs

To characterize cell populations targeted by nnAbs and bnAbs in HIV-1-infected primary CD4+ T cells, we used multiparametric flow cytometry to simultaneously probe CD4 and viral proteins expression. Activated primary CD4+ T cells were mock-infected or infected with the transmitted-founder virus CH077 (CH077TF). Two days post-infection, cells were stained with a panel of nnAbs and bnAbs, followed by appropriate secondary Abs. Cells were then stained for cell-surface CD4 prior to staining for intracellular HIV-1 p24. Given that the *nef* gene is abundantly expressed during the early phase of the HIV-1 replication cycle (63, 64), we also evaluated the expression of this accessory protein by intracellular staining as previously described (61). The different cell populations were gated based on cell-surface CD4 and intracellular p24 detection as shown in Figure 1A. As expected, uninfected CD4^+^p24^−^ cells were not recognized by bnAbs of various specificities (PGT126, PG9, 3BNC117, PGT151 and 2G12) (Figure 1B-D). Interestingly, this population was efficiently recognized by CD4i nnAbs targeting the coreceptor-binding site (17b) or the gp120 inner domain (A32, C11) as well as a pool of purified immunoglobulins from PLWH (HIV-IG). However, the CD4^+^p24^−^ cells were not recognized by the CD4i gp41-specific nnAbs F240. The absence of binding by bnAbs and F240 suggested that these cells were likely coated with shed gp120 rather than presenting CD4-bound cell-surface Env trimer. This is in line with previous work showing that uninfected CD4+ T cells expose CD4i epitopes on their cell surface after interacting with gp120 shed from productively-infected cells within the same culture (33, 65, 66). Indeed, introduction of a CD4bs (D368R) mutation into CH077TF that prevents Env-CD4 interaction (17, 23, 67), abrogated the recognition of the CD4^+^p24^−^ population by all gp120-specific nnAbs tested (Figure S1A-B).

**Figure 1.**
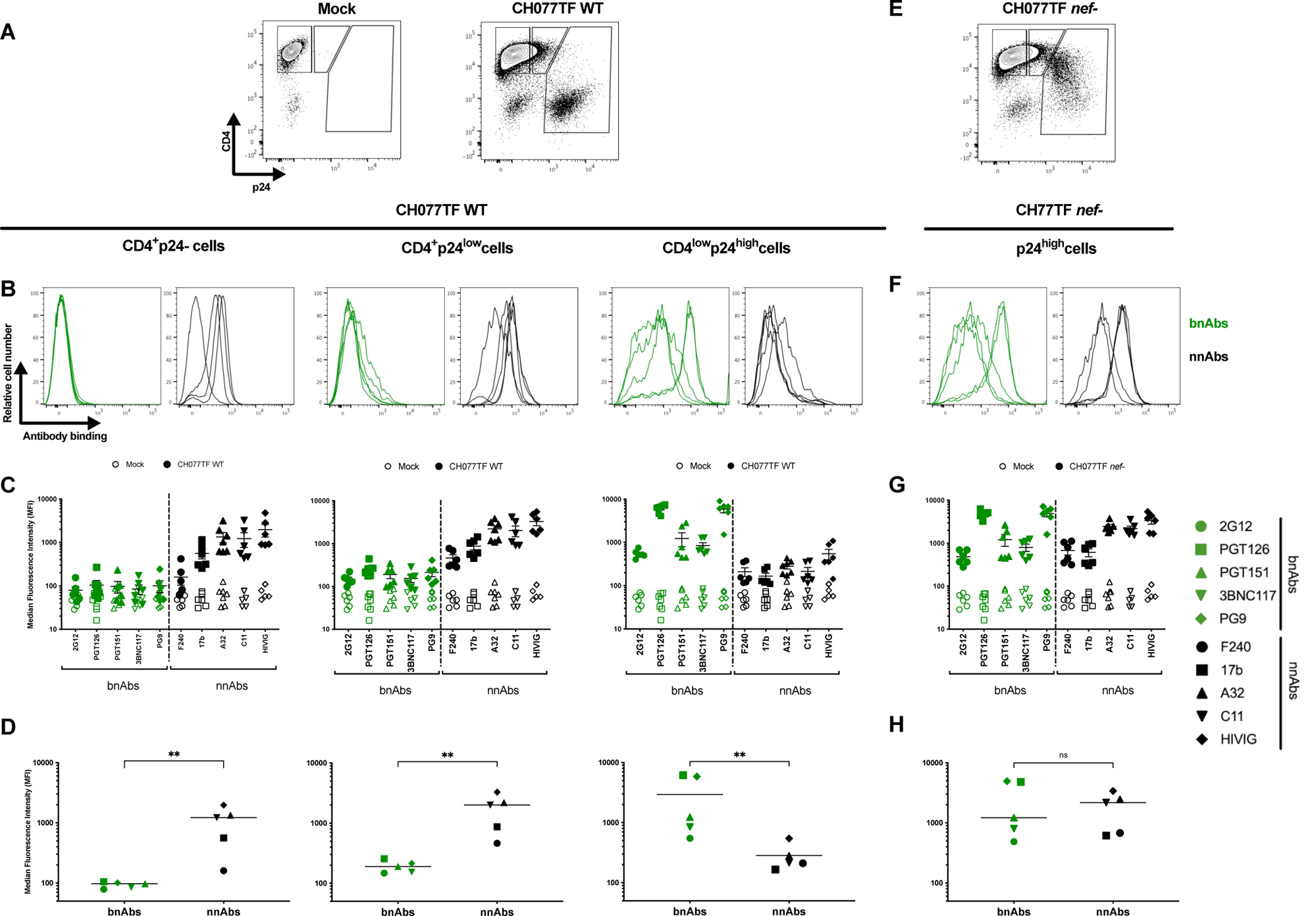
Recognition of HIV-1-infected primary CD4+ T cells by bnAbs and nnAbs. Primary CD4+ T cells, mock-infected or infected with the transmitted-founder virus CH077, either wild-type (WT) or defective for Nef expression (*nef*-) were stained with a panel of bnAbs and nnAbs, followed with appropriate secondary Abs. Cells were then stained for cell-surface CD4 prior detection of intracellular HIV-1 p24. (A,E) Example of flow cytometry gating strategy based on cell-surface CD4 and intracellular p24 detection. (B,F) Histograms depicting representative staining with bnAbs (Green) and nnAbs (Black) (C,G). Graphs shown represent the median fluorescence intensities (MFI) obtained for at least 6 independent staining with the different mAbs. Error bars indicate means ± standard errors of the means. (D,H) Graphs shown represent the mean MFI obtained with each mAb. Statistical significance was tested using Mann-Whitney U test (** p<0.01, ns: non-significant).

In addition to productively infected CD4+ T cells, which efficiently downregulated CD4 (CD4^low^p24^high^), we identified a subset of CD4^+^p24^low^ cells, as previously reported (51, 58, 68–72). This CD4^+^p24^low^ subset was efficiently recognized both by the gp120-specific nnAbs and HIV-IG, but was resistant to bnAbs binding. This population was also recognized by the gp41-specific F240 nnAb, suggesting the presence of trimeric Env bound to CD4. However, recognition of the CD4^+^p24^low^ cells by CD4i nnAbs was substantially reduced upon the introduction of the D368R mutation (Figure S1A-B). This mutation also specifically reduced the proportion of CD4^+^p24^low^ cells (Figure S1C). These findings, along with the observation that these cells do not express Nef (Figure 2) suggest that CD4^+^p24^low^ cells are not productively infected, but contain CD4-Env complexes on their surface resulting from the binding of shed gp120 and/or viral particles.

**Figure 2.**
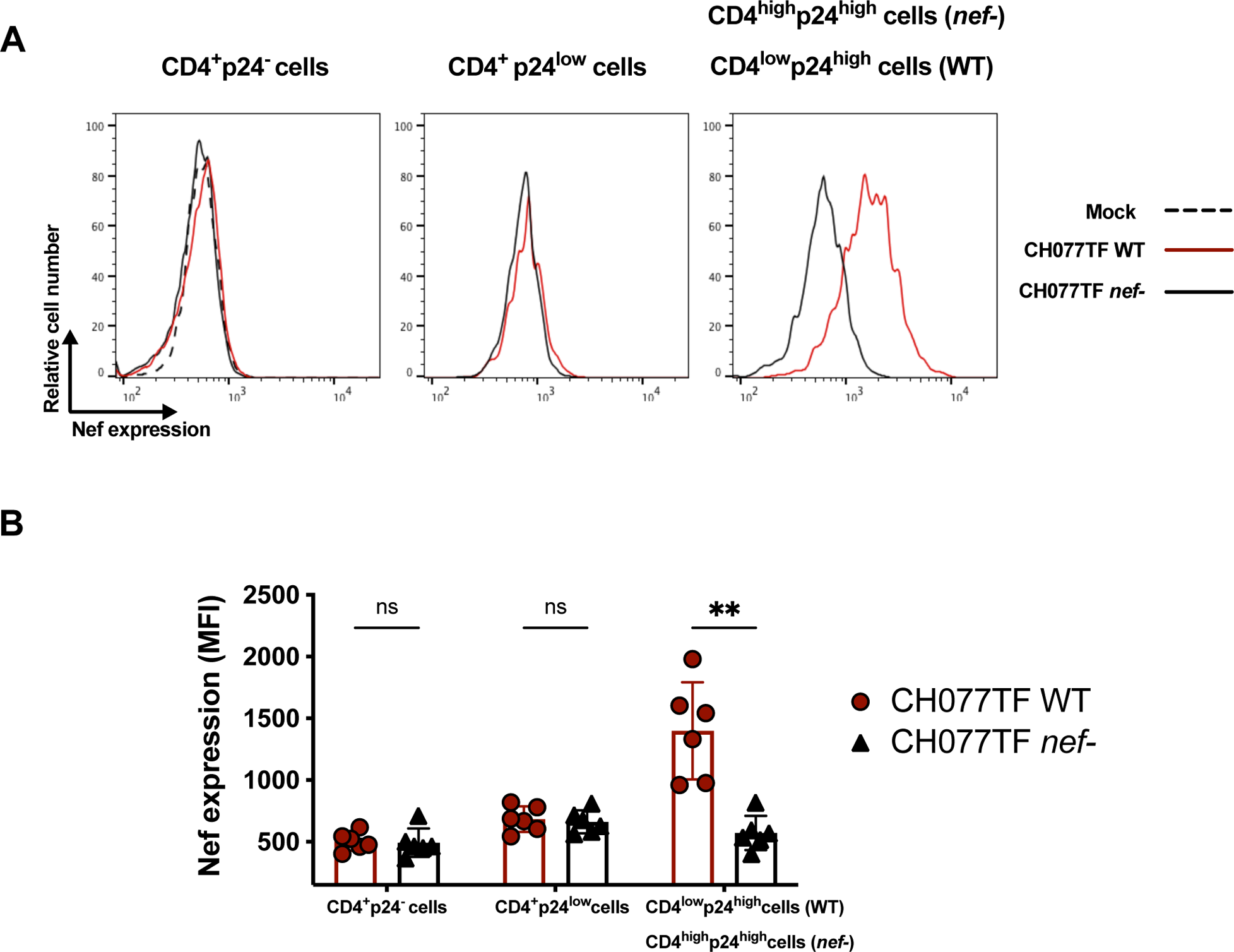
Nef is expressed in HIV-1-infected cells undergoing CD4 downregulation and expressing high levels of p24. Primary CD4+ T cells, mock-infected or infected with the transmitted-founder virus CH77, either WT or defective for Nef expression (Nef-) were stained for cell-surface CD4 prior detection of intracellular HIV-1 p24 and Nef expression. (A) Histograms depicting representative intracellular Nef staining when gating on CD4^+^p24^−^, CD4^+^p24^low^ or CD4^low^p24^high^ cells. In the context of cells infected with CH77TF Nef-, in the absence of Nef-mediated CD4 downmodulation, the p24^high^ remained CD4^high^ (CD4^high^p24^high^). (B) Quantification of the median fluorescence intensities (MFI) obtained for 6 independent experiments. Error bars indicate means ± standard errors of the means. Statistical significance was tested using multipe Mann-Withey tests with a Holm-Sidak post-test (** p<0.01, ns: non-significant).

As expected, cells that expressed high level of p24 (CD4^low^p24^high^) were poorly recognized by nnAbs. We reasoned that this was because they efficiently downregulated cell-surface CD4, which precludes premature Env triggering, and thus prevents the exposure of normally occluded epitopes (23, 26, 33). CD4^low^p24^high^ cells also expressed Nef (Figure 2) and were efficiently recognized by bnAbs, known to preferentially recognize Env in its “closed” conformation (Figure 1B-D). However, when we used a *nef*-defective virus these cells were efficiently recognized by nnAbs (Figure 1E-H), in agreement with previous observations (26, 33, 37, 39, 61, 73, 74).

To test whether our findings extend to IMCs widely used in the ADCC field (28, 35, 53, 55, 68, 72, 75–77), we infected primary CD4+ T cells with with viruses produced from a pNL4.3 backbone expressing different Envs, including BaL and NL4.3. As shown in Figure S2A-B, the patterns of bnAbs and nnAbs binding were very similar to those observed for CH077TF. The CD4^+^p24^−^ and CD4^+^p24^low^ cell populations were preferentially recognized by the nnAbs, while the CD4^low^p24^high^ cells were efficiently targeted by the bnAbs. Interestingly, despite similar levels of productive infection (CD4^low^p24^high^, Figure S2C), an enrichement of the CD4^+^p24^low^ population was detected in the context of infection with the NL4.3 and BaL Envs as compared to Envs from primary viruses (CH058TF, CH040TF, SF162 and YU2) (Figure S2D). The reasons for this are unclear but may be due to greater gp120 shedding of the tier 1 NL4.3 and BaL Envs.

Taken together, our results indicate that cells recognized by nnAbs express high levels of CD4, are either p24^low^ or p24^−^, and are negative for Nef expression. In contrast, bnAbs recognize cells that efficiently downregulate CD4 and express high level of p24 and Nef proteins.

### e*nv* mRNA is predominantly detected in cells that already downregulated CD4

To better understand the underlying mechanisms behind the differential recognition of infected cells by bNAbs versus nnAbs, we used a previously described RNA-flow cytometric fluorescence *in situ* hybridization (RNA Flow-FISH) method (78–80). This method identifies productively-infected cells by detecting cellular HIV-1 mRNA by using *in situ* RNA hybridization and intracellular Ab staining for the HIV-1 p24 protein. In the context of these experiments, *env* and *nef* mRNA probes were used to identify productively-infected CD4+ T cells. Briefly, primary CD4 T cells were mock-infected or infected with the CH077TF IMC. Two days post-infection, cells were stained for surface CD4 before fixation and permeabilization to allow detection of the HIV-1 p24 antigen and HIV-1 mRNAs. Cell populations were first defined based on their cell-surface CD4 and intracellular p24 co-expression, as presented in Figure 3A. Productively-infected cells were identified as *nef* mRNA+/*env* mRNA+.The vast majority of cells with detectable cell-surface CD4 (CD4^+^p24^−^ or CD4^+^p24^low^) were negative for HIV-1 mRNA (Figure 3B-C), while cells that efficiently downmodulated CD4 (CD4^low^p24^low^ and CD4^low^p24^high^) were enriched for *nef* and *env* mRNA transcripts (Figure 3B-C; Figure S3). When gating on productively-infected cells based on *nef* and *env* mRNA detection, we confirmed that the CD4^low^p24^high^ cells represent the major source of productively-infected cells (Figure 3D-E). These results show that *env* mRNA is predominantly expressed by HIV-1-infected cells that already downregulated CD4 (CD4^low^p24^low^ and CD4^low^p24^high^) (Figure S3). This analysis also captured stages of infection (CD4^low^p24^low^) where CD4 is already downmodulated while *env* mRNA expression intensifies, suggesting that CD4 downmodulation precedes *env* mRNA expression.

**Figure 3.**
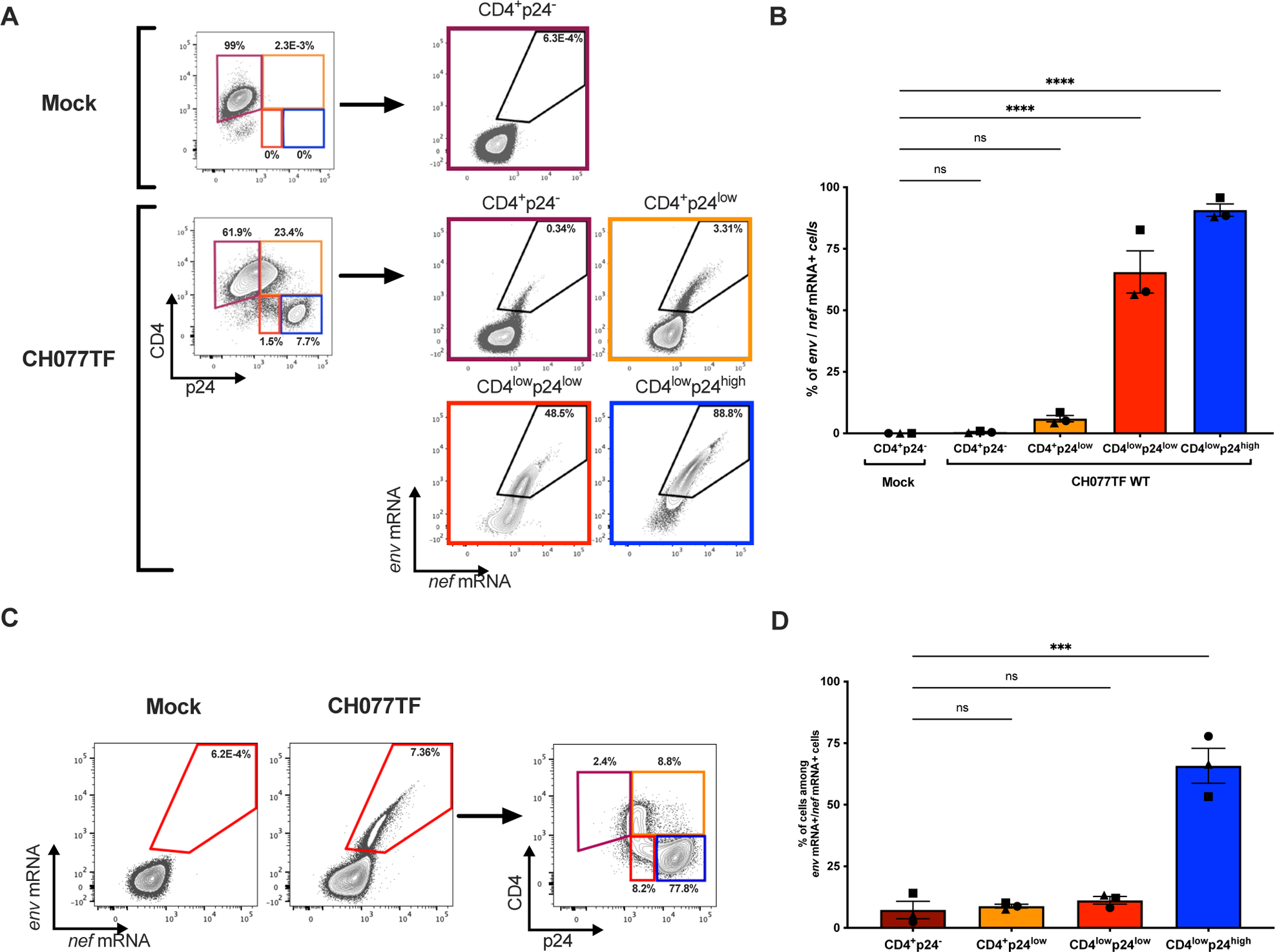
HIV-1 late transcripts are mostly detected among cells that downregulated CD4. Purified primary CD4+ T cells, mock-infected or infected with the transmitted-founder virus CH077 WT were stained for cell-surface CD4 prior detection of intracellular HIV-1 p24 and *env* mRNA and *nef* mRNA by RNA-flow FISH. (A) Representative example of flow cytometry gating strategy based on cell-surface CD4 and intracellular p24 detection. (B) Representative example of RNA-flow FISH detection of *env* and *nef* mRNA among the different cell populations. (C) Quantification of the percentage of *env* mRNA+ *nef* mRNA+ cells detected among the different cell populations in three different donors. (D) Alternatively, productively-infected cells were first identified based on *env* and *nef* mRNA detection (E) Quantification of the percentage of CD4^+^p24^−^, CD4^+^p24^low^, CD4^low^p24^low^ and CD4^low^p24^high^ cells among the *env* and *nef* mRNA+ cells with three different donors. Statistical significance was tested using one-way ANOVA with a Holm-Sidak post-test (****p<0.0001, ns: non-significant).

### Cells targeted by A32 are negative for HIV-1 mRNA

We next combined flow cytometry and RNA flow-FISH methods to compare the capacity of nnAbs and bnAbs to bind productively-infected cells. Mock-infected or CH077TF-infected primary CD4 T cells were first stained with the nnAb A32 or the bNAb PGT126. Cells where then stained for cell-surface CD4 detection, intracellular p24 and HIV-1 mRNAs. Productively-infected cells were identified based on the simultaneous detection of *nef* and *env* mRNA transcripts (Figure 4A). As shown in Figure 4B-C, productively-infected cells (*env*/*nef* mRNA+ cells) were recognized by PGT126, but not by A32. In contrast, *env*/*nef* mRNA-cells were targeted by A32, but not by PGT126. Similar results were obtained when cells were classified by Ab recognition (Figure 4D-F; Figure S4), with the majority of PGT126+ cells co-expressing HIV-1 mRNA. In contrast, cells recognized by A32 were negative for *env* and *nef* mRNA (Figure 4E-F; Figure S4). These results indicate that cells targeted by A32 do not express *env* mRNA (Figure S4), are not productively infected and thus are likely coated with gp120 and/or viral particles.

**Figure 4.**
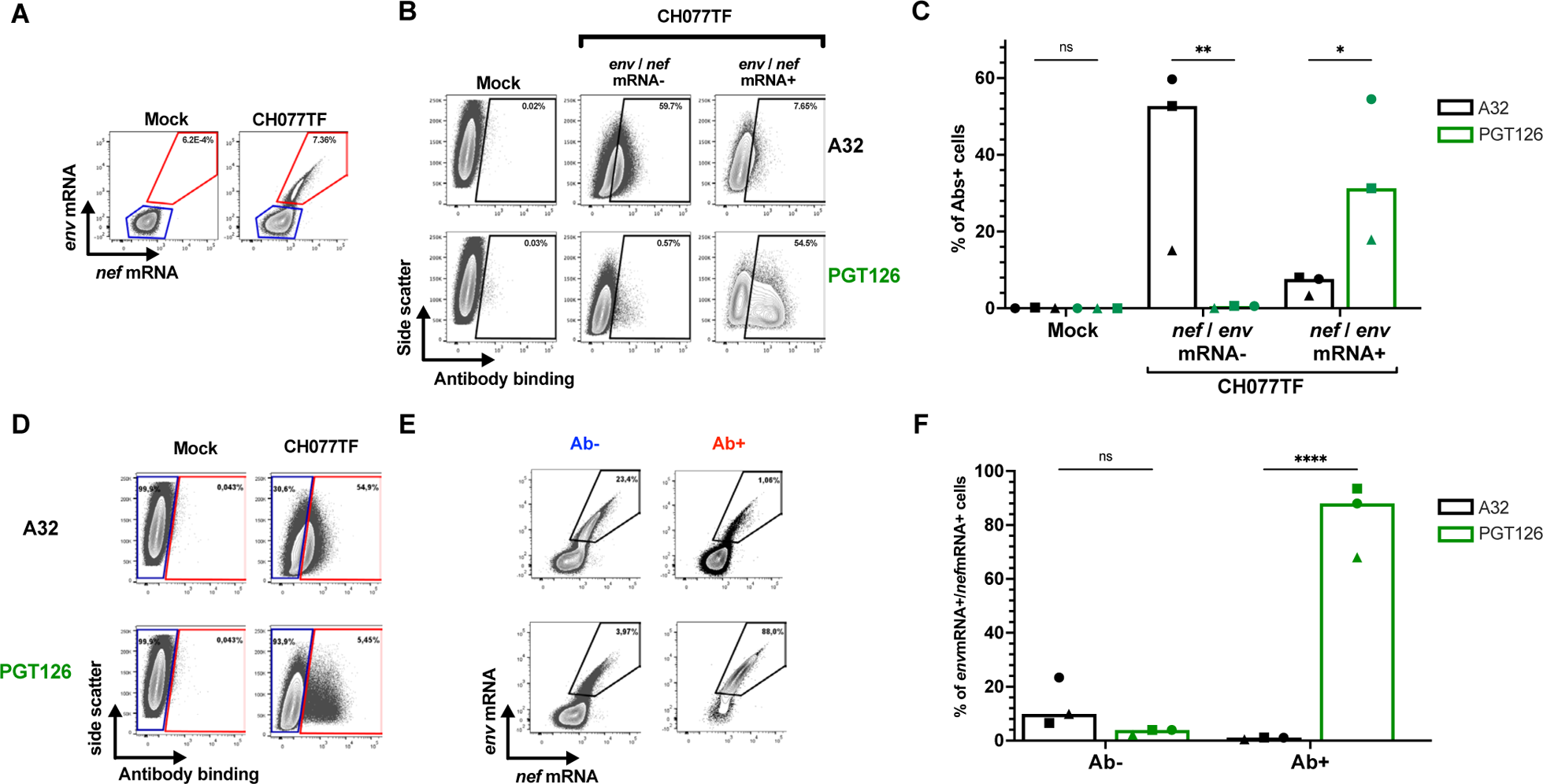
Productively-infected cells are resistant to recognition by A32. Purified primary CD4+ T cells, mock-infected or infected with the transmitted-founder virus CH077 WT were stained with A32 or PGT126, followed with appropriate secondary Abs. Cells were then stained for cell-surface CD4 prior detection of intracellular HIV-1 p24 and *Env* mRNA and *nef* mRNA by RNA-flow FISH. (A-C) In a first analysis, HIV-infected cells were identified, then A32 and PGT126 binding was evaluated. (A) Example of RNA-flow FISH gating strategy based on *env* and *nef* mRNA detection. (B) Example of antibody binding among the *env/nef* mRNA- and *env/nef* mRNA+ cell population. (C) Quantification of the percentage of cells recognized by either A32 or PGT126 among the *env/nef* mRNA-and *env/nef* mRNA+ cell population with three different donors. (D-F) In a second alternative analysis, Ab-binding cells were first identied, and the HIV-infection status was then evaluated. (D) Example of flow cytometry gating strategy based on A32 or PGT126 binding. (E) Example of *env/nef* mRNA detection among the cells recognized (Ab+) or not (Ab-) by indicated mAbs. (F) Quantification of the percentage of *env/nef* mRNA+ cells among the cells recognized (Ab+) or not (Ab-) by indicated mAbs with three donors. Statistical significance was tested using a two-way ANOVA with a Holm-Sidak post-test (* p<0.05,** p<0.01, **** p<0.0001, ns: non-significant).

### *Ex vivo* expanded CD4+ T cells isolated from PLWH are resistant to ADCC responses mediated by nnAbs

Our results indicated that productively infected cells are principally CD4^low^p24^high^, express *nef* and *env* mRNA and are not recognized by nnAbs. To confirm these findings with primary clinical samples, we expanded CD4+ T cells from PLWH. Briefly, CD4+ T cells were isolated from 6 chronically-infected individuals and activated *ex vivo* using PHA-L/IL-2 (24, 38, 40). CD4+ T cells from people without HIV were used as controls. Viral replication was monitored over time by intracellular p24 staining (Figure 5A). Upon expansion, CD4+ T cells were stained with a panel of bnAbs and nnAb, followed by the appropriate secondary Abs. Cells were stained for cell-surface CD4 prior to the detection of intracellular HIV-1 p24 and Nef proteins. Consistent with the results obtained with IMC infections (Figure 2), productively-infected CD4^low^p24^high^ cells were the only ones which were also positive for the Nef protein (Figure S5). These cells were efficiently recognized by bnAbs and largely resistant to nnAbs binding (Figure 5C,D, E). In contrast, nnAbs mainly recognized CD4+ cells that were either p24^−^ or p24^low^ as well as negative for Nef expression (Figure 5C, D, E; Figure S5). To evaluate the susceptibility of productively-infected cells to ADCC responses mediated by bnAbs and nnAbs, expanded endogenously-infected cells were used as target cells and autologous peripheral blood mononuclear cells (PBMCs) as effectors using a FACS-based ADCC assay (Figure 6). Consistent with antibody binding, productively-infected CD4^low^p24^high^ cells were resistant to ADCC mediated by nnAbs, but sensitive to those mediated by bnAbs.

**Figure 5.**
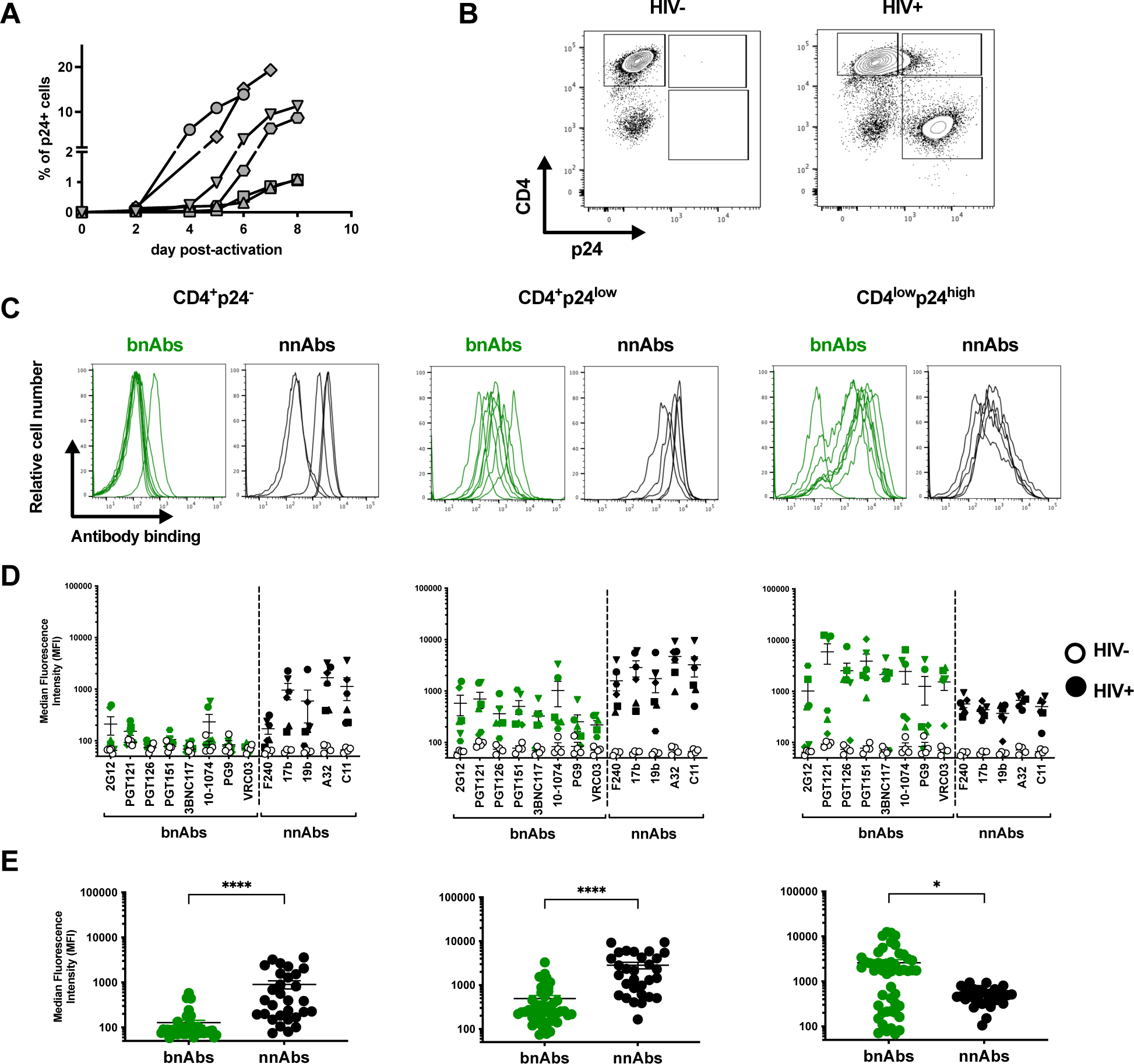
*Ex vivo* expanded CD4+ T cells isolated from PLWH are preferentially targeted by bNAbs. *Ex vivo* expanded CD4+ T cells from 6 PLWH were stained with bnAbs and nnAbs +/− CD4mc, followed by appropriate secondary Abs. Cells were then stained for surface CD4 prior to detection of intracellular HIV-1 p24. (A) Percentage of p24+ upon activation overtime. (B) Example of flow cytometry gating based on CD4 and p24 detection. (C) Histograms depicting representative staining with bnAbs (Green) and nnAbs (Black). (D) Median fluorescence intensities (MFI) obtained with 6 HIV+ individuals. (E) Graphs shown represent the mean MFI obtained with each mAb. Each symbol represent a diffirent HIV+ donor. Statistical significance was tested using Mann-Whitney U test (* p<0.05, **** p<0.0001).

**Figure 6.**
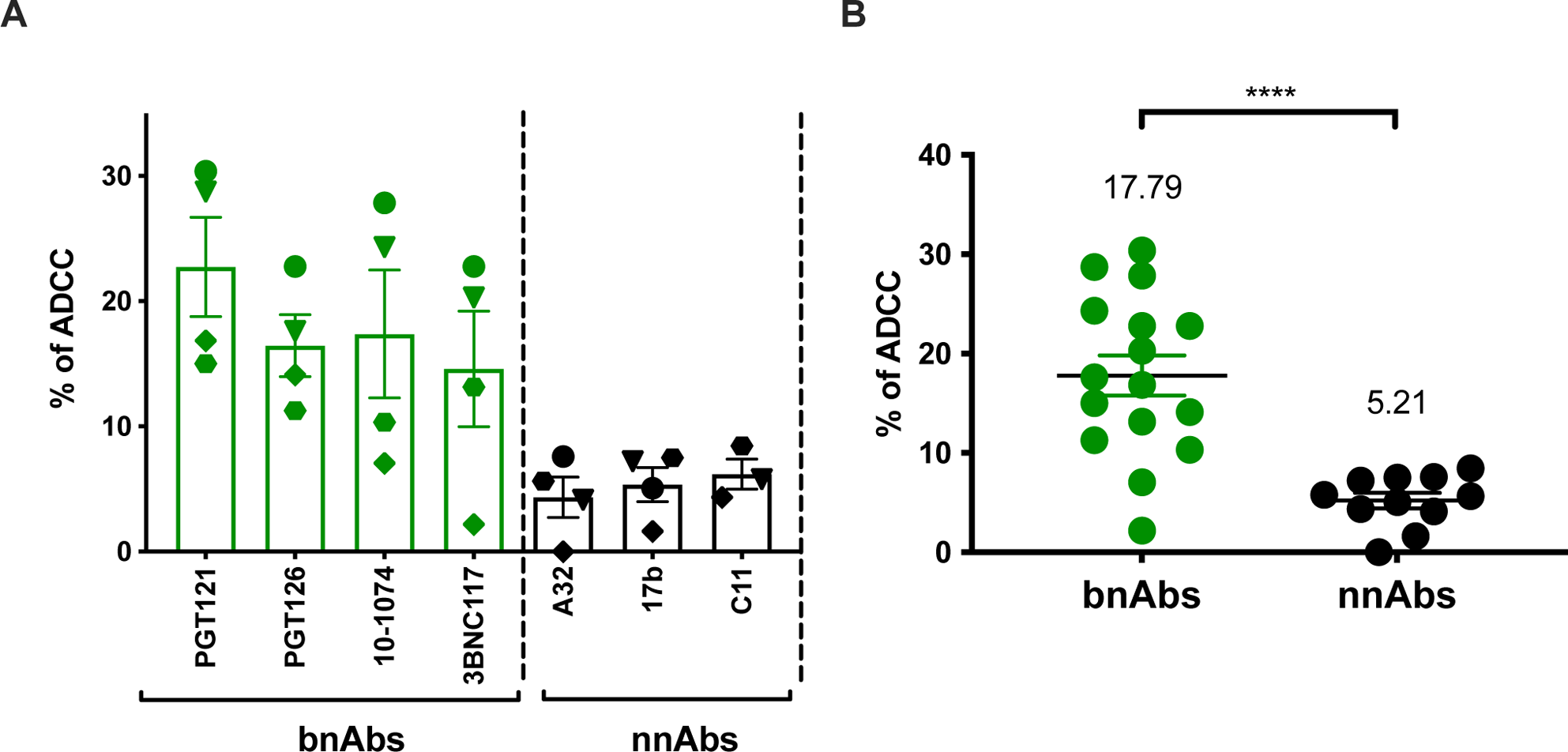
*Ex vivo* expanded CD4+ T cells isolated from PLWH are resistant to ADCC mediated by nnAbs. *Ex vivo* expanded CD4 T cells from PLWH were used as target cells, while autologous PBMC were used as effector cells in our FACS-based ADCC assays. (A) Graph shown represent the percentage of ADCC against the CD4^low^p24^high^ cells with the single mAbs and (B) nnAbs vs bnAbs. (A) Each symbol represent a diffirent HIV+ donor. Statistical significance was tested using unpaired t test (**** p<0.0001).

### A32 does not affect HIV-1 replication or the size of HIV-1 reservoir *in vivo*

Our results indicate that nnAbs, such as A32, target nonproductively-infected CD4+ T cells. It has been suggested that these cells could be in a very early stage of infection, during viral entry, before viral gene expression (81). Specifically, it has previously been shown that non-neutralizing Env epitopes, such as that targeted by A32, become transiently exposed during viral entry and could therefore represent a suitable target for ADCC at this stage (81). We hypothesized that if this was the case, then this cell population should be eliminated by A32 and therefore decrease HIV-1 replication *in vivo*. To evaluate this possibility, we tested whether A32 affected viral replication in humanized mice (hu-mice). Briefly, NOD.Cg-*Prkdc^scid^* IL2rg^−/−^ Tg(Hu-IL15) (NSG-15) hu-mice engrafted with human peripheral blood lymphocytes (hu-PBL) were infected with the primary isolate HIV-1JRCSF (Figure 7A). This hu-mice model was previously shown to support HIV-1 replication and antibody Fc-effector function *in vivo* (24, 82). Infected humice received nnAb A32 administered subcutaneously (S.C.) at day 6 and 9 post-infection (Figure 7A). As controls, infected mice were also treated with the bnAbs 3BNC117 or its Fc gamma receptor (FcγR) null binding variant (GRLR) (83). Hu-mice were monitored for plasma viral loads (PVLs) and peripheral CD4+ T cells overtime (Figure 7B-C). In the absence of antibody treatment, mice became viremic, reaching an average PVL of 2.2×10^7^ copies/mL at day 11. As previously reported (41, 82), viral replication was associated with a loss of peripheral CD4+ T cells. Treatment with 3BNC117 WT significantly reduced viral replication and partially restored CD4+ T cell levels in peripheral blood and several tissues, while A32 failed to do so (Figure 7B,C,D). Interestingly, A32 treatment further reduced CD4+ T cell levels in tissues relative to mock-treated mice (Figure 7D), consistent with its capacity to recognize and eliminate uninfected bystander cells coated with soluble gp120 via ADCC (33, 65, 66). This reduction in CD4 T cells count was particularly significant when CD4+ T cell levels where considered across all tissues in mice treated with A32 nnAb alone (Figure S6). Finally, treatment with 3BNC117 WT led to a significant reduction of the HIV-1 reservoir in multiple tissues, a phenotype not observed with A32 and not fully achieved by its FcγR null binding variant (3BNC117 GRLR, Figure 7E). These results are in line with previous work demonstrating that bnAbs require Fc effector functions for *in vivo* activity (83). These results indicate that A32 does not reduce HIV-1 replication or the size of the reservoir in hu-mice.

**Figure 7.**
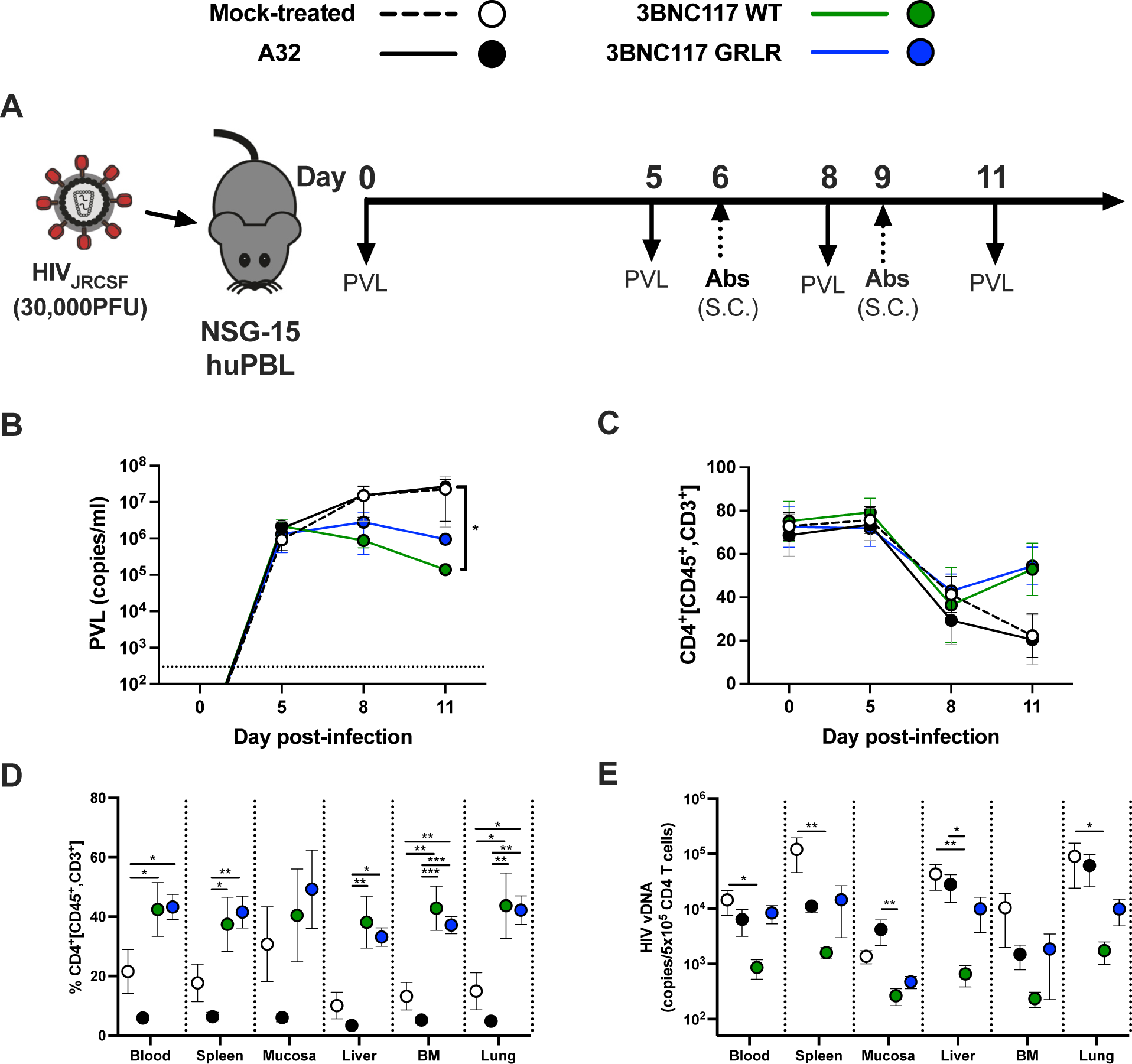
A32 nnAb does not impact viral replication or the size of the reservoir *in vivo*. (A) Experimental outline. NSG-15-Hu-PBL mice were infected with HIV-1 JRCSF intraperitoneally. At day 6 and 9 post infection, mice were administered 1.5 mg of A32 or 3BNC117 (WT or GRLR) mAb subcutaneously (S.C.). (B) Mice were bled routinely for plasma viral load (PVL) and flow cytometry analysis. PVL levels were measured by quantitative real-time PCR (limit of detection = 300 copies/mL, dotted line). (C) Percentage of CD4+ T cells in peripheral blood was evaluated by flow cytometry. At least six mice were used for each treatment. (D-E) Tissues of JRCSF-infected NSG-15 hu-PBL mice, treated or not with A32 or 3BNC117 (WT or GRLR) were harvested at day 11. (D) Percentage of CD4+ T cells was evaluated by flow cytometry (E) CD4+ T cells were isolated for real-time PCR analysis of HIV DNA. Each dot represents the mean values +/− SEM. S.C., subcutaneous; I.P., intraperitoneal; BM, bone marrow; mock-treated = no antibody. Statistical significance was tested using one way ANOVA with a Holm-Sidak post-test or a Kruskal-Wallis test with a Dunn’s post-test (* p<0.05,** p<0.01,*** p<0.001)

## DISCUSSION

ADCC represents an effective immune response involved in the clearance of virally-infected cells. This adaptive immune response relies on the capacity of Abs to act as a bridge between infected cells and effector cells. While the Abs recognize infected cells through binding of surface Env via their Fab domain, their Fc domain allows the recruitment and the stimulation of FcγR bearing effector cells, leading to the killing of infected cells. ADCC-mediating Abs therefore represent attractive therapeutics for HIV cure strategies. However, the ability of Abs to recognize productively-infected cells and to mediate ADCC responses depends on Env conformation (73). While bnAbs target epitopes on the prefusion “closed” Env trimer, nnAbs recognize conserved epitopes that are normally occluded but are exposed when Env interacts with CD4 and adopt an “open” conformation (29, 32–36). Here, we provide evidence that *env* mRNA expression is mainly restricted to cells that already downmodulated cell-surface CD4 (Figures 3 and S3). This prevents CD4i Env epitopes exposure on productively-infected cells, thus contributing to their resistance to CD4i nnAbs.

Non-neutralizing Abs, such as those targeting the CoRBS and the inner domain of gp120 are elicited during natural infection due to exposure to “viral debris” acting as immunodominant decoy (23, 25, 28, 34, 84, 85). It is therefore not surprising that HIV-1 has evolved mechanisms to prevent surface Env-CD4 interaction to avoid Env recognition by these nnAbs which have potent Fc-effector functions. HIV-1 utilizes Nef, Vpu and Env to downmodulate CD4 from the surface of infected cells (86). Nef is expressed at high levels early during infection from a multi-spliced transcript (63, 64) and downregulates CD4 by enhancing its internalization and lysosome degradation (87–91). Vpu and Env are expressed from a Rev-depended single spliced bicistronic mRNA later during the viral life cycle (92). Both Vpu and Env interfere with the transport of newly synthesized CD4 to the cell surface (93–95). Using RNA-Flow FISH methods, we show that *env* mRNA is almost exclusively detected in cells positive for *nef* mRNA that efficiently downregulated CD4. The majority of cells expressing HIV-1 mRNA substantially downregulated surface CD4 and express high levels of p24 and Nef proteins (CD4^low^p24^high^ cells). We also captured stage of infection (CD4^low^p24^low^) where CD4 is already downmodulated while *env* mRNA and p24 are not fully expressed. This suggest that Nef-mediated CD4 downmodulation precedes Env expression (Figure 3 and S3). Accordingly, productively-infected cells failed to expose CD4i epitopes, explaining their resistance to ADCC mediated by nnAbs, while showing susceptibility to ADCC mediated by bNabs (Figure 1, 4, 5 and 6).

Our findings are consistent with a growing number of studies demonstrating the importance of bnAbs in mediating ADCC against cells productively infected with primary HIV-1 isolates (32, 33, 35). nnAbs recognize productively-infected cells only when CD4 is not properly downregulated, as is the case in infections with Nef defective virus (Figure 1). This is also the case when *nef*-deficient IMCs are used for ADCC detection (28, 49–60, 69, 71, 76). Most of these IMCs express the Renilla luciferase (LucR) reporter gene upstream of the *nef* sequence and use a T2A ribosome-skipping peptide to promote Nef expression (62). Despite documented evidence of reduced Nef expression (39, 61, 62), these IMCs continue to be used in the ADCC field (51, 52, 54, 55, 57, 58, 96). While some studies have suggested that Vpu can compensate for the absence of Nef (51, 52), our findings refute this. Given its early expression and ability to target cell surface expressed CD4 molecules, Nef plays the most prominent role in CD4 downmodulation (61, 86, 97). In absence of Nef expression, we find that Vpu is not sufficient to downregulate cell-surface CD4, leading to CD4i epitope exposure and efficient nnAbs binding (Figure 1). Other studies reporting Fc-effector functions of nnAbs employed assays unable to differentiate the ADCC responses directed against HIV-1-infected cells versus uninfected bystander cells (45, 49, 76, 98, 99). The presence of uninfected bystander cells coated with gp120 shed from productively-infected cells impacts ADCC measurement by introducing a significant bias toward CD4i nnAbs (33), which is also the case when ADCC assays rely solely on target cells coated with gp120 or inactivated virions (33, 50, 60, 71, 72, 100, 101). Utilization of such assays, as well as *nef*-defective IMCs, contribute to the propagation of a misleading concept that nnAbs can effectively mediate ADCC against HIV-1-infected cells. nnAbs such as A32 not only fail to eliminate HIV-1-infected cells, but also have potentially detrimental effects by accelerating the elimination of uninfected bystander cells (33, 66), as shown in tissues of HIV-1-infected humanized mice (Figure 7; Figure S6). In this context, the absence of an antiviral effect of A32 in hu-mice is not surprising. Studies in non-human primate showed that elicitation of A32-like Abs by gp120 immunization or passive administration of A32 failed to confer protection against simian-human immunodeficiency virus challenges (77, 102). Similarly, a combination of anti-CoRBS and anti-cluster A (A32) nnAbs proved ineffective in delaying viral rebound after ART interruption in humanized mouse model supporting Fc effector functions (41).

The inability of A32 to recognize productively-infected cells, to influence viral replication and reduce the size of reservoir in hu-mice is a function of its epitope, which is occluded in the unliganded trimer. To our knowledge, exposure of this epitope at the surface of productively infected cells is possible in only two ways: either by membrane-bound CD4 (26) or the combination of potent small CD4-mimetic compounds (CD4mc) and anti-CoRBS Abs (27, 41). Of note, the cocktail of A32, 17b (CoRBS Ab) and CD4mc was reported to significantly reduce the size of the reservoir in hu-mice (41).

In conclusion, we show that *env* mRNA is almost exclusively expressed by productively-infected cells that downregulated cell-surface CD4. This suggests that CD4 downmodulation precedes Env expression, thus preventing exposure of vulnerable CD4-induced Env epitopes and evading ADCC mediated by nnAbs. These results must be taken into account when considering the use of nnAbs for preventative or cure strategies.

## ACKNOWLEDGMENTS

The authors thank the CRCHUM BSL3 and Flow Cytometry Platforms for technical assistance, and Mario Legault from the FRQS AIDS and Infectious Diseases network for cohort coordination and clinical samples. The graphical abstract was created using BioRender.com. We thank the following collaborators for kindly providing plasmids to produce antibodies: James Robinson (Tulane University) for A32 and C11, the NIH AIDS Reagent Program for F240, 2G12 and 17b, John Mascola (Vaccine Research Center, NIAID) for VRC03, the International AIDS Vaccine Initiative (IAVI) for PG9, PGT121, PGT126 and PGT151 and Michel Nussenzweig (Rockefeller University) for 3BNC117 and 10-1074.

This study was supported by grants from the National Institutes of Health to A.F. (R01 AI148379 and R01 AI150322, R01 AI129769, R01AI176531) and P.K. (R01AI145164). Support for this work was also provided by P01 GM56550/AI150471 to A.F., a Canadian Institutes of Health Research (CIHR) Team grant #422148 to D.E.K, P.K. and A.F., a CIHR foundation grant #352417 to A.F. and a Canada Foundation for Innovation (CFI) grant #41027 to D.E.K. and A.F. This work was partially supported by UM1AI144462 (CHAVD) (D.E.K), UM1AI164562 (ERASE) to A.F., and P.K., and UM1AI164559 (HOPE) to P.K., and by an American Foundation for AIDS Research (amfAR) Mathilde Krim Fellowship in Basic Biomedical Research (109720-63-RKRL) to J.R. A.F. was supported by a Canada Research Chair on Retroviral Entry #RCHS0235 950-232424. B.H.H. is supported by R01 AI162646, UM1AI144371 and UM1AI164570. J.P. was supported by a CIHR doctoral fellowship. M.B. was supported by a FRQS master’s fellowship. G.B.B was supported by a FRQS and a CIHR doctoral fellowship. FK is supported by the German Research Foundation (DFG, CRC 1279) and an ERC Advanced grant (Traitor viruses). The funders had no role in study design, data collection and analysis, decision to publish, or preparation of the manuscript.

## AUTHOR CONTRIBUTIONS

**Conceptualization:** J.R., G.S, L.Z, P.K., D.E.K and A.F.; **Methodology:** J.R, G.S, J.P., L.Z..,M.D., P.K., D.E.K. and A.F.; **Investigation:** J.R, G.S., L.Z., L.M., M.B., G.B.B., D.C., Y.S., H.K., J.P., H.M., C.B., and G.G.D.; **Resources:** J.R., G.S., J.P., L.Z., L.M., M.B., G.B.B., D.C., F.K., B.H.H., P.K., D.E.K. and A.F.; **Formal Analysis:** J.R., G.S. and L.Z.; **Visualization:** J.R., G.S and L.Z. **Supervision:** M.D., P.K., D.E.K. and A.F.; **Funding acquisition:** J.R. P.K., D.E.K. and A.F.; **Writing – original draft:** J.R., B.H.H., and A.F. **Writing – review & editing:** J.R., G.S., L.Z., J.P., L.M., M.B., G.B.B., D.C., F.K., B.H.H.,Y.S., H.K., H.M., C.B., G.G.D., M.D., P.K., D.E.K. and A.F.

## METHODS

### Ethics Statement

Written informed consent was obtained from all study participants and research adhered to the ethical guidelines of CRCHUM and was reviewed and approved by the CRCHUM institutional review board (ethics committee, approval number MP-02-2024-11734). Research adhered to the standards indicated by the Declaration of Helsinki. All participants were adult and provided informed written consent prior to enrolment in accordance with Institutional Review Board approval.

### Cell lines and primary cells

293T human embryonic kidney cells (obtained from ATCC) were maintained at 37°C under 5% CO2 in Dulbecco’s Modified Eagle Medium (DMEM) (Wisent, St. Bruno, QC, Canada), supplemented with 5% fetal bovine serum (FBS) (VWR, Radnor, PA, USA) and 100 U/mL penicillin/streptomycin (Wisent). Human peripheral blood mononuclear cells (PBMCs) from HIV-negative individuals and HIV-positive individuals obtained by leukapheresis and Ficoll-Paque density gradient isolation were cryopreserved in liquid nitrogen until further use. CD4+ T lymphocytes were purified from resting PBMCs by negative selection using immunomagnetic beads per the manufacturer’s instructions (StemCell Technologies, Vancouver, BC) and were activated with phytohemagglutinin-L (10 µg/mL) for 48 h and then maintained in RPMI 1640 (Thermo Fisher Scientific, Waltham, MA, USA) complete medium supplemented with rIL-2 (100 U/mL).

### Antibody production

FreeStyle 293F cells (Thermo Fisher Scientific) were grown in FreeStyle 293F medium (Thermo Fisher Scientific) to a density of 1 × 10^6^ cells/mL at 37°C with 8% CO2 with regular agitation (150 rpm). Cells were transfected with plasmids expressing the light and heavy chains of each mAb using ExpiFectamine 293 transfection reagent, as directed by the manufacturer (Thermo Fisher Scientific). One week later, the cells were pelleted and discarded. The supernatants were filtered (0.22-μm-pore-size filter), and antibodies were purified by protein A affinity columns, as directed by the manufacturer (Cytiva, Marlborough, MA, USA).

### HIV-1 studies in humanized mice

NSG-15 mice with expression of the human IL15 gene in the NOD/ShiLtJ background were purchased from the Jackson Laboratory (Bar Harbor, ME, USA). The mice were bred and maintained under specific pathogen-free conditions. All animal studies were performed with authorization from Institutional Animal Care and Use Committees (IACUC) of Yale University. NSG-15-Hu-PBL mice were engrafted as described (41). Briefly, 3.5 × 10^6^ PBMCs, purified by Ficoll density gradient centrifugation of healthy donor blood buffy coats, obtained from the New York Blood Bank) were injected IP in a 200-µL volume into 6- to 8-week-old NSG-15 mice, using a 1-cm^3^ syringe and 25-gauge needle. Cell engraftment was tested 15 days post-transplant. 100 µL of blood was collected by retroorbital bleeding. PBMCs were isolated by Ficoll density gradient centrifugation; stained with fluorescently-labelled anti-human CD45 (BD Biosciences, Cat#: 555485), CD3 (Biolegend, Cat#: 300424), CD4 (Biolegend, Cat#: 317432), CD8 (BD Biosciences, Cat#: 561617) and CD56 (Biolegend, Cat#: 362508) antibodies and analyzed by flow cytometry to confirm engraftment. Humanized mice were intraperitoneally challenged with 30,000 PFU of HIV-1JRCSF. Infection profile was analyzed routinely by retro-orbital bleeding and flow cytometric analysis of peripheral blood for human immune cells and PVL analysis. For flow cytometry, 100 μl of blood was collected by retro-orbital bleeding at each time point. PBMCs were isolated by Ficoll density gradient centrifugation and cells were stained with fluor-conjugated antibodies as detailed above. PVL were measured at day 5, 8 and 11 post-infection, while HIV-1 reservoirs were measured at day 11 post-infected as previously described (41).

### Plasmids and proviral constructs

Transmitted/Founder (T/F) infectious molecular clone (IMC) of patient CH077 was inferred and constructed as previously described (103). The generation of *nef*-defective CH077TF was previously described and consists in the introduction of premature stop codons in the *nef* reading frame using site-directed mutagenesis protocol (104). The CH077TF D368R was also generated by site-directed mutagenesis as previously described (42). Proviral constructs comprising an HIV-1 NL4.3-based isogenic backbone engineered for the insertion of heterologous *env* strain sequences and expression in *cis* of full length Env (Env-IMCs), were previously described (105). The Env-IMCs utilized in the present study are those encoding Env form BaL (pNL-B.BaL.ecto), CH040 (pNL-B.CH040.ecto), CH058 (pNL-B.CH058.ecto), SF162 (pNL-B.SF162.ecto) or YU2 (pNL-B.YU-2.ecto). The proviral plasmid pNL4.3 was used as control (106). The vesicular stomatitis virus G (VSV-G)-encoding plasmid was previously described (107).

### Viral production, infections and *ex vivo* amplification

For *in vitro* infection, vesicular stomatitis virus G (VSV-G)-pseudotyped HIV-1 viruses were produced by co-transfection of 293T cells with an HIV-1 proviral construct and a VSV-G-encoding vector using the calcium phosphate method. Two days post-transfection, cell supernatants were harvested, clarified by low-speed centrifugation (300 × g for 5 min), and concentrated by ultracentrifugation at 4°C (100,605 × g for 1 h) over a 20% sucrose cushion. Pellets were resuspended in fresh RPMI, and aliquots were stored at −80°C until use. Viruses were then used to infect activated primary CD4+ T cells from healthy HIV-1 negative donors by spin infection at 800 × *g* for 1 h in 96-well plates at 25 °C. Viral preparations were titrated directly on primary CD4+ T cells to achieve similar levels of infection among the different IMCs tested (around 10% of p24+ cells). To expand endogenously infected CD4+ T cells, primary CD4+ T cells obtained from PLWH were isolated from PBMCs by negative selection. Purified CD4+ T cells were activated with PHA-L at 10 μg/mL for 48 h and then cultured for at least 6 days in RPMI 1640 complete medium supplemented with rIL-2 (100 U/ml) to reach greater than 10% infection for the ADCC assay.

### Antibodies

The following anti-Env Abs were used to stained HIV-1-infected primary CD4+ T cells: anti-gp41 ID F240; anti-cluster A A32, C11; anti-co-receptor binding site 17b; anti-gp120 outer domain 2G12, anti-CD4 binding site VRC03, 3BNC117; anti-V3 glycan PGT121, PGT126, 10-1074; anti-V2 apex PG9; anti-gp120-gp41 interface PGT151. The HIV-IG polyclonal antibody consists of anti-HIV immunoglobulins purified from a pool of plasma from HIV+ asymptomatic donors (NIH AIDS Reagent Program). Goat anti-human IgG (H+L) (Thermo Fisher Scientific) pre-coupled to Alexa Fluor 647 were used as secondary antibodies in flow cytometry experiments. Rabbit antisera raised against Nef (NIH AIDS Reagent Program) was used as primary antibodies in intracellular staining. BrillantViolet 421 (BV421)-conjugated donkey anti-rabbit antibodies (Biolegend) was used as secondary antibodies to detect Nef antisera binding by flow cytometry. FITC or PE-conjugated Mouse anti-human CD4 (clone OKT4; Biolegend) were used for cell-surface staining of HIV-1-infected primary CD4+ T cells, while PE or FITC-conjugated Mouse anti-HIV-1 p24 (clone KC57; Beckman coulter) were used for intracellular staining.

### Flow cytometry analysis of cell-surface staining

Cell surface staining was performed at 48h post-infection. Mock-infected or HIV-1-infected primary CD4+ T cells were incubated for 30 min at 37°C with anti-Env mAbs (5 µg/mL) or HIV-IG (50 µg/mL). Cells were then washed once with PBS and stained with the appropriate Alexa Fluor 647-conjugated secondary antibody (2 µg/mL) for 20 min at room temperature. Cells were then stained with FITC- or PE-conjugated Mouse anti-CD4 Abs. After two PBS washs, cells were fixed in a 2% PBS-formaldehyde solution. Infected cells were then permeabilized using the Cytofix/Cytoperm Fixation/ Permeabilization Kit (BD Biosciences, Mississauga, ON, Canada) and stained intracellularly using PE or FITC-conjugated mouse anti-p24 mAb (clone KC57; Beckman Coulter, Brea, CA, USA; 1:100 dilution). The percentage of infected cells (p24^+^) was determined by gating on the living cell population according to a viability dye staining (Aqua Vivid; Thermo Fisher Scientific). Alternatively, cells were stained intracellularly with rabbit antisera raised against Nef (1:1000) followed by BV421-conjugated anti-rabbit secondary antibody. Samples were acquired on an LSR II cytometer (BD Biosciences), and data analysis was performed using FlowJo v10.5.3 (Tree Star, Ashland, OR, USA).

### Antibody-dependant cellular cytotoxicity (ADCC) assay

Measurement of ADCC using a fluorescence-activated cell sorting (FACS)-based infected cell elimination (ICE) assay was performed at 48 h post-infection. Briefly, HIV-1-infected primary CD4+ T cells were stained with AquaVivid viability dye and cell proliferation dye eFluor670 (Thermo Fisher Scientific) and used as target cells. Cryopreserved autologous PBMC effectors cells, stained with cell proliferation dye eFluor450 (Thermo Fisher Scientific), were added at an effector: target ratio of 10:1 in 96-well V-bottom plates (Corning, Corning, NY). Anti-Env mAbs (5 µg/mL) was added to appropriate wells and cells were incubated for 5 min at room temperature. The plates were subsequently centrifuged for 1 min at 300 × g, and incubated at 37 °C, 5 % CO_2_ for 5 h. before being stained with FITC- or PE-conjugated Mouse anti-CD4 Abs. After one PBS wash, cells were fixed in a 2% PBS-formaldehyde solution. Infected cells were then permeabilized using the Cytofix/Cytoperm Fixation/ Permeabilization Kit (BD Biosciences, Mississauga, ON, Canada) and stained intracellularly using PE or FITC-conjugated mouse anti-p24 mAb (clone KC57; Beckman Coulter, Brea, CA, USA; 1:100 dilution) Productively-infected cells were identified based on p24 and CD4 detection as described above. Samples were acquired on an LSR II cytometer (BD Biosciences) and data analysis was performed using FlowJo v10.5.3 (Tree Star). The percentage of ADCC was calculated with the following formula: [(% of CD4^+^p24^high^ cells in Targets plus Effectors) − (% of CD4^+^p24^high^ cells in Targets plus Effectors plus plasma or mAbs) / (% of CD4^+^p24^high^ cells in Targets) × 100] by gating on infected lived target cells.

### RNAflow-FISH analysis

All buffers and fixation reagents were provided with the kit, with the exception of flow cytometry staining buffer (2% FCS/PBS). The HIV-1 RNAflow-FISH assay was performed as previously described and as per manufacturer’s instructions (78–80). Briefly, cells were harvested 48h post-infection and stained with the anti-Env mAbs A32 or PGT126 as described above. Cells were then stained with Fixable Viability Dye (20 min, 4°C, Fixable LiveDead, eBioscience) and then with a mix containing a brilliant stain buffer (BD Biosciences) and the surface markers for CD4+ T cells detection (CD3 and CD4) and CD8/NK/B cells and macrophages exclusions (CD8, CD56, CD19, CD16) (30 min, 4°C). Samples were fixed, permeabilized with buffers provided by the manufacturer, and labeled intracellularly for the structural HIV-1 p24 protein with the anti-p24 clone KC57 antibody (30 min RT followed by 30 min 4°C, Beckman Coulter). HIV-1 RNA probing was performed using the PrimeFlow RNA Assay (ThermoFisher). HIV-1 RNA were labeled using HIV-1 *env*RNA (Thermofisher; catalog number VF6-6000978) and HIV-1 *nef*RNA (Thermofisher; catalog number (VF4-6000647) probe sets. HIV-1 *env*RNA was designed based on CH077TF full env sequence, whereas HIV-1 *nef*RNA was based on consensus B HIV-1 sequence. Each probeset allows the hybridization of specific complementary branched DNA nanostructure with different excitation/emission spectra. The probeset were diluted 1:5 in diluent and hybridized to the target mRNAs for 2 hr at 40°C. Samples were washed to remove excess probes and stored overnight in the presence of RNAsin. Signal amplification was achieved by performing sequential hybridization with DNA branches (i.e., Pre-Amplifier and Amplifier) The first DNA branch in the Pre-Amplifier Mix was added at a 1:1 ratio and was allowed to hybridize for 1.5 h at 40°C. Then the second DNA branch in the Amplifier Mix was added and hybridized for 1.5 h at 40°C (78–80). Amplified mRNAs were labeled with fluorescently tagged probes allowing hybridization for 1 hr at 40°C. Samples were acquired on an LSRFortessa (BD Biosciences) and analyzed using FlowJo (BD, V10.7.0). Unspecific binding of the fluorescent labeled branched probe in the multiplex kit can lead to a low level of false-positive background noise, which, if present, is detected across all the four channels corresponding to the types of labeled probes (AF488, AF594, AF647, AF750). To decrease background noise, we thus left the AF594 channel vacant and excluded false-positive events based on fluorescence in this channel before further gating. Gates were set on the HIV-uninfected donor control, or unstimulated control where appropriate.

## QUANTIFICATION AND STATISTICAL ANALYSIS

Statistics were analyzed using GraphPad Prism version 9.1.0 (GraphPad, San Diego, CA, USA). Every data set was tested for statistical normality and this information was used to apply the appropriate (parametric or nonparametric) statistical test. P values <0.05 were considered significant; significance values are indicated as * P<0.05, ** P<0.01, *** P<0.001, **** P<0.0001.

### Data availability

The published article includes all datasets generated and analyzed for this study. Further information and requests for resources and reagents should be directed to and will be fulfilled by the Lead Contact Author. All unique reagents generated in this study are available from the Lead Contact with a completed Materials Transfer Agreement.

## Authorship note

JR, GS and LZ contributed equally to this work and PK, DEK and AF contributed equally as co-senior authors. AF is the lead contact

## SUPPLEMENTAL FIGURE LEGENDS

**Figure S1.**
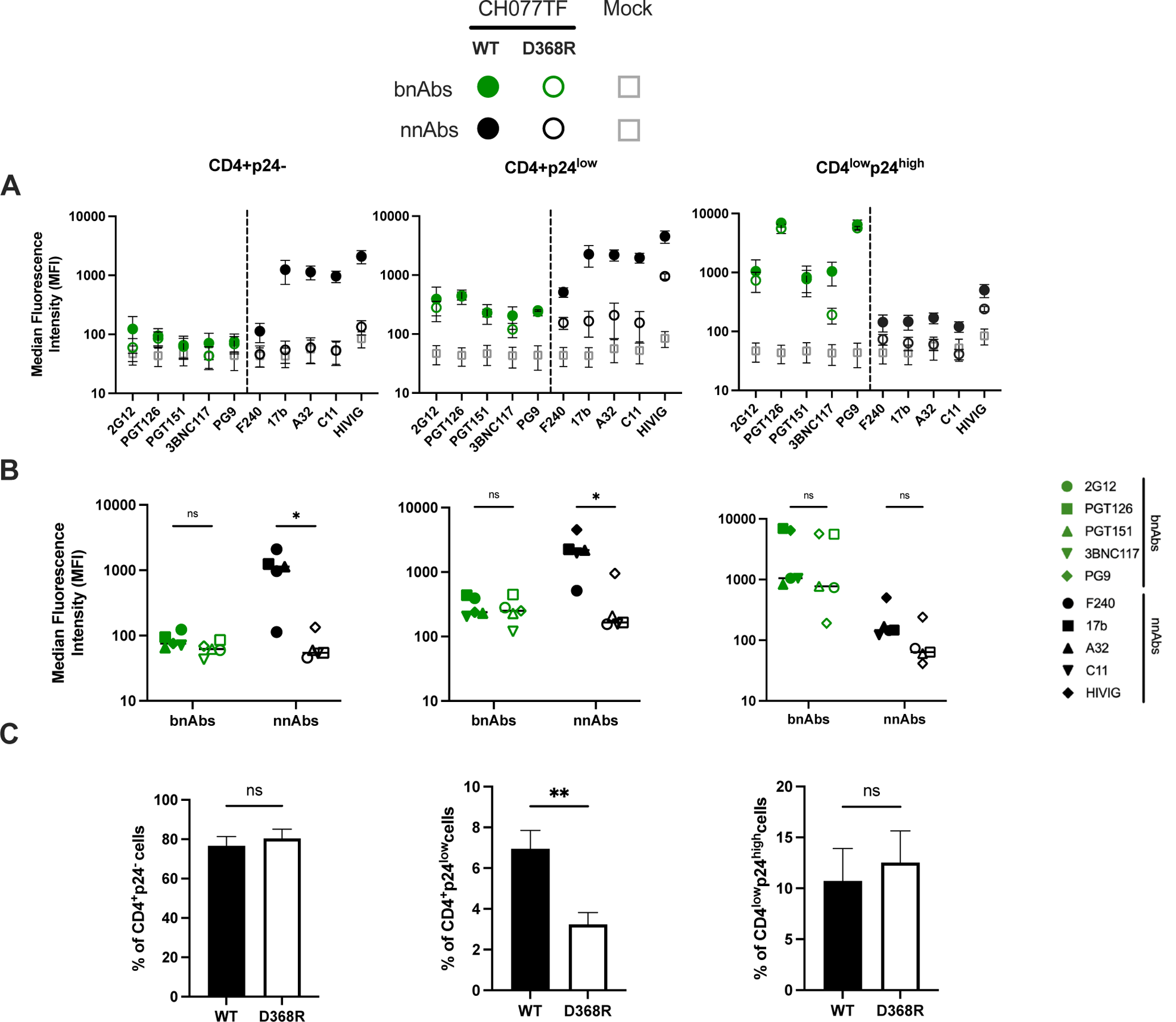
Impact of the D368R mutation on nnAbs and bnAbs binding. Primary CD4+ T cells, mock-infected or infected with the transmitted-founder virus CH077, either expressing the wild-type (WT) or D368R Env (D368R) were stained with a panel of bnAbs and nnAbs, followed with appropriate secondary Abs. Cells were then stained for cell-surface CD4 prior detection of intracellular HIV-1 p24. (A) Graphs shown represent the median fluorescence intensities (MFI) obtained for at least 3 independent staining with the different mAbs. Error bars indicate means ± standard errors of the means. (B) Graphs shown represent the mean MFI obtained with each mAbs. (C) Percentage of the different cell populations at 48h post-infection. Statistical significance was tested using Mann-Whitney U test (** p<0.01, ns: non-significant).

**Figure S2.**
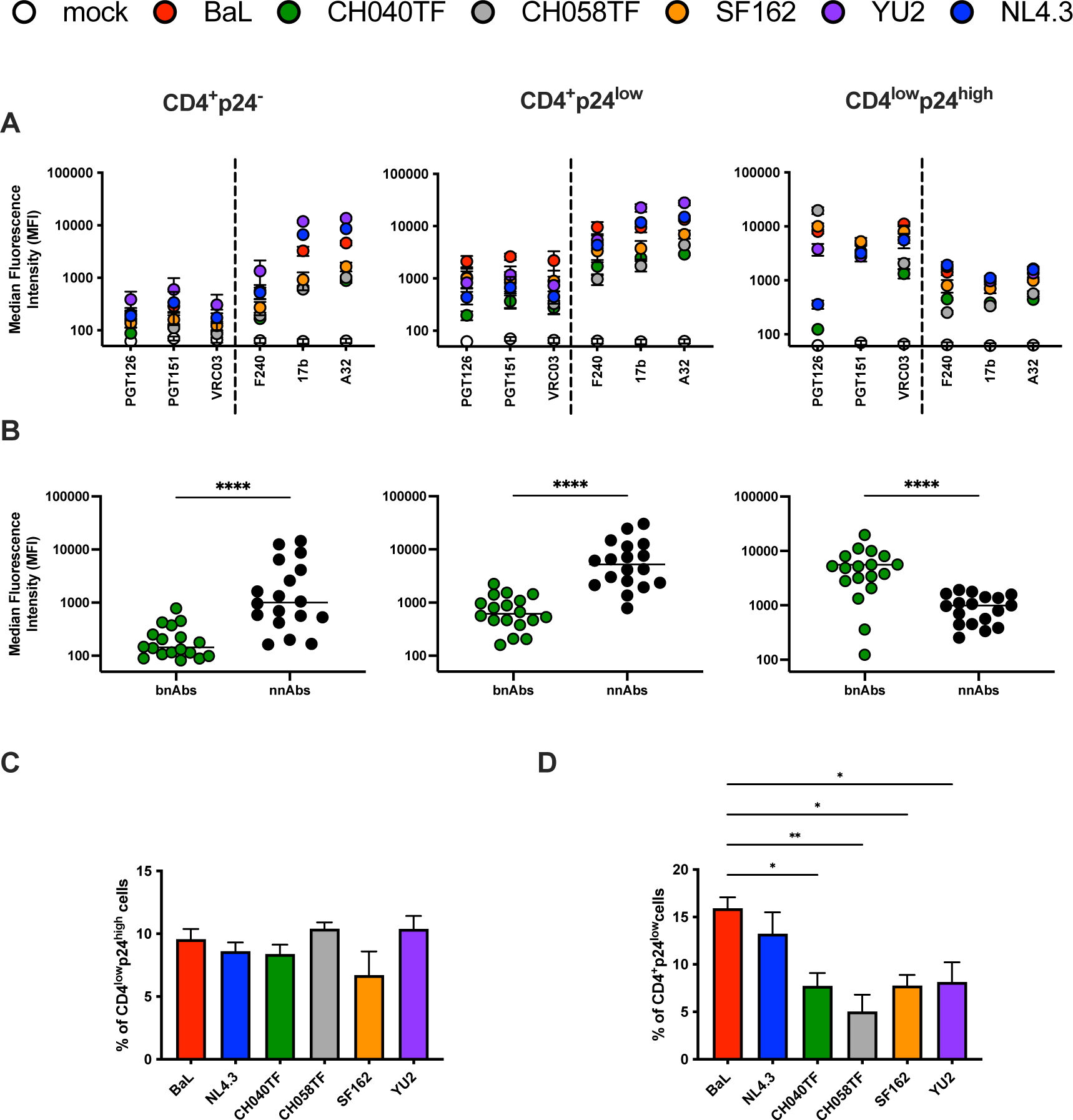
Recognition of cells infected with HIV-1 constructs expressing lab-adapted and primary Envs by bnAbs and nnAbs. Primary CD4+ T cells, mock-infected or infected with NL4.3 infectious molecular clones (IMC) expressing Env from lab-adapted (BaL, NL4.3) or primary (CH040TF, CH058, YU2, SF162) viruses were stained with a panel of bnAbs and nnAbs, followed with appropriate secondary Abs. Cells were then stained for cell-surface CD4 prior detection of intracellular HIV-1 p24. (A) Graphs shown represent the median fluorescence intensities (MFI) obtained for at least 3 independent staining with the different mAbs. Error bars indicate means ± standard errors of the means. (B) Graphs shown represent the mean MFI obtained with each mAbs for each HIV-1 IMC. (C-D) Percentage of the (C) CD4^low^p24^high^ and (D) CD4^+^p24^low^ populations at 48h post-infection. Statistical significance was tested Mann-Whitney U test (* p<0.05, ** p<0.01, **** p<0.0001).

**Figure S3.**
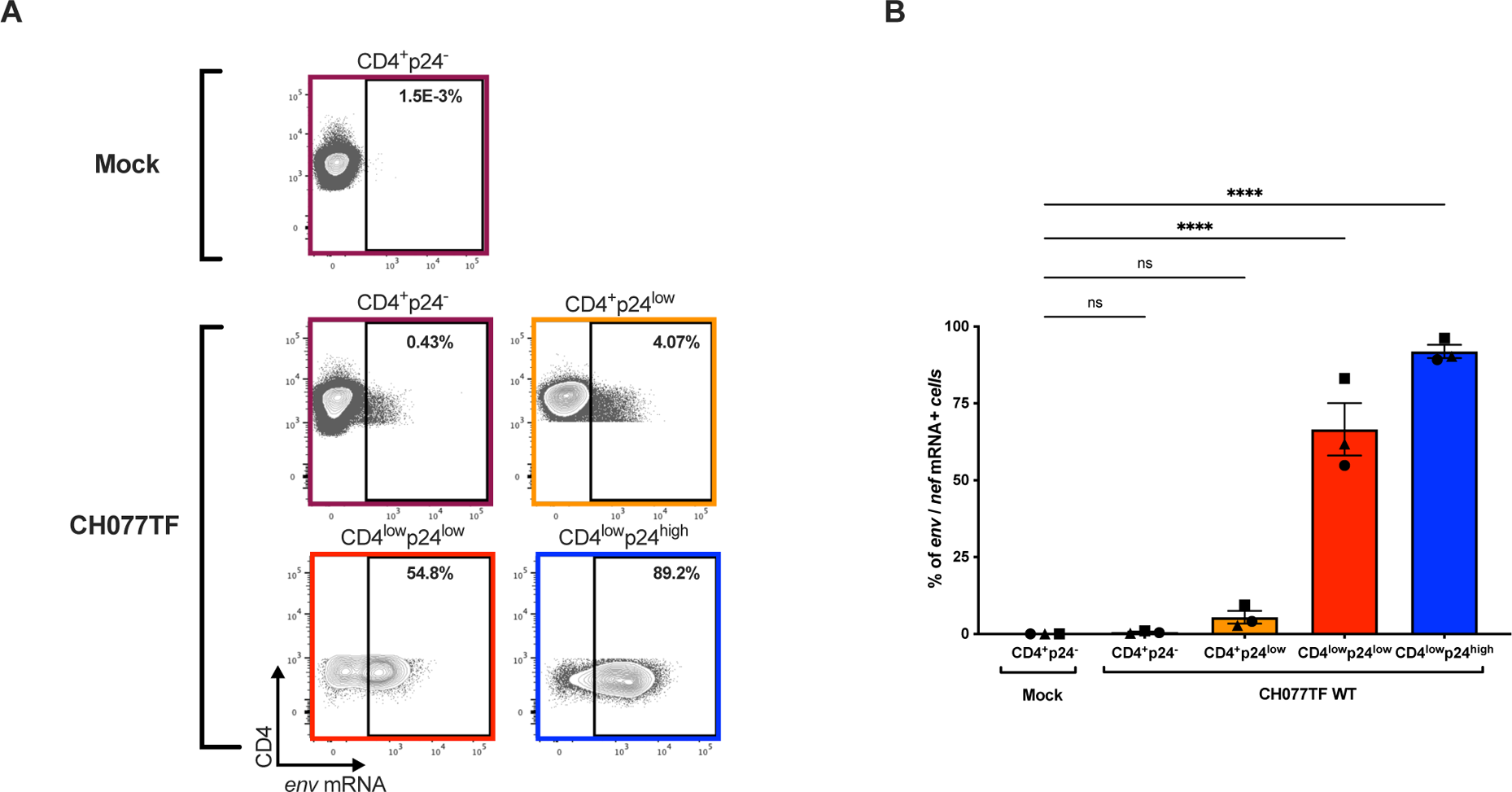
CD4 downregulation precedes Env expression. Primary CD4+ T cells, mock-infected or infected with the transmitted-founder virus CH077 WT were stained for cell-surface CD4 prior detection of intracellular HIV-1 p24 and *Env* mRNA and *nef* mRNA by RNA-flow FISH. (A) Example of RNA-flow FISH detection of *env* mRNA among the CD4^+^p24^−^, CD4^+^p24^low^, CD4^low^ p24^low^ or CD4^low^ p24^high^ cell populations. (C) Quantification of the percentage of *env* mRNA+ cells detected among the different cell population with three different donors. Statistical significance was tested using one way ANOVA with a Holm-Sidak post-test (* p<0.05, ** p<0.01, ns: non-significant).

**Figure S4.**
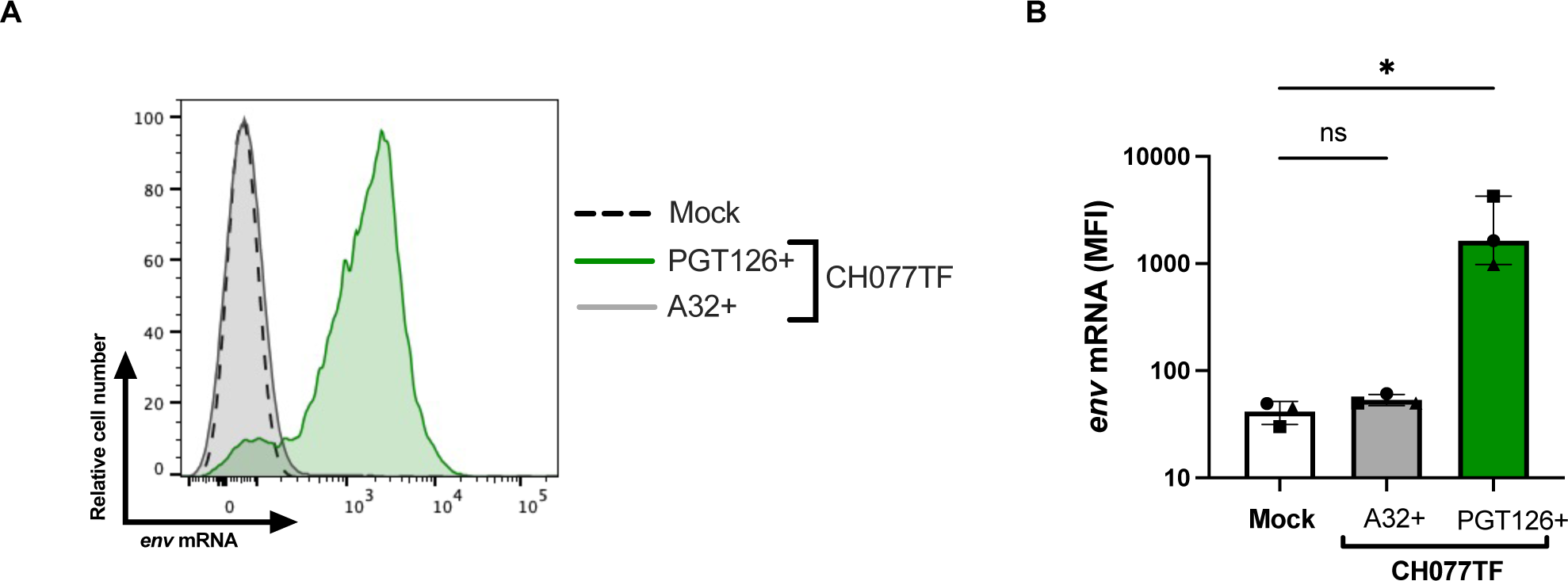
Cells targeted by A32 are *env* mRNA negative. Primary CD4+ T cells, mock-infected or infected with the transmitted-founder virus CH77 WT were stained with A32 or PGT126, followed with appropriate secondary Abs. Cells were then stained for cell-surface CD4 prior detection of intracellular HIV-1 p24 and *Env* mRNA and *nef* mRNA by RNA-flow FISH. (A) Histograms depicting representative *env* mRNA detection on A32+ or PGT126+ cells. (B) Median fluorescence intensities (MFI) of *env* mRNA obtained with 3 different donors. Statistical significance was tested using a Kruskal-Wallis test with a Dunn’s post-test. (* p<0.05, ns: non-significant)

**Figure S5.**
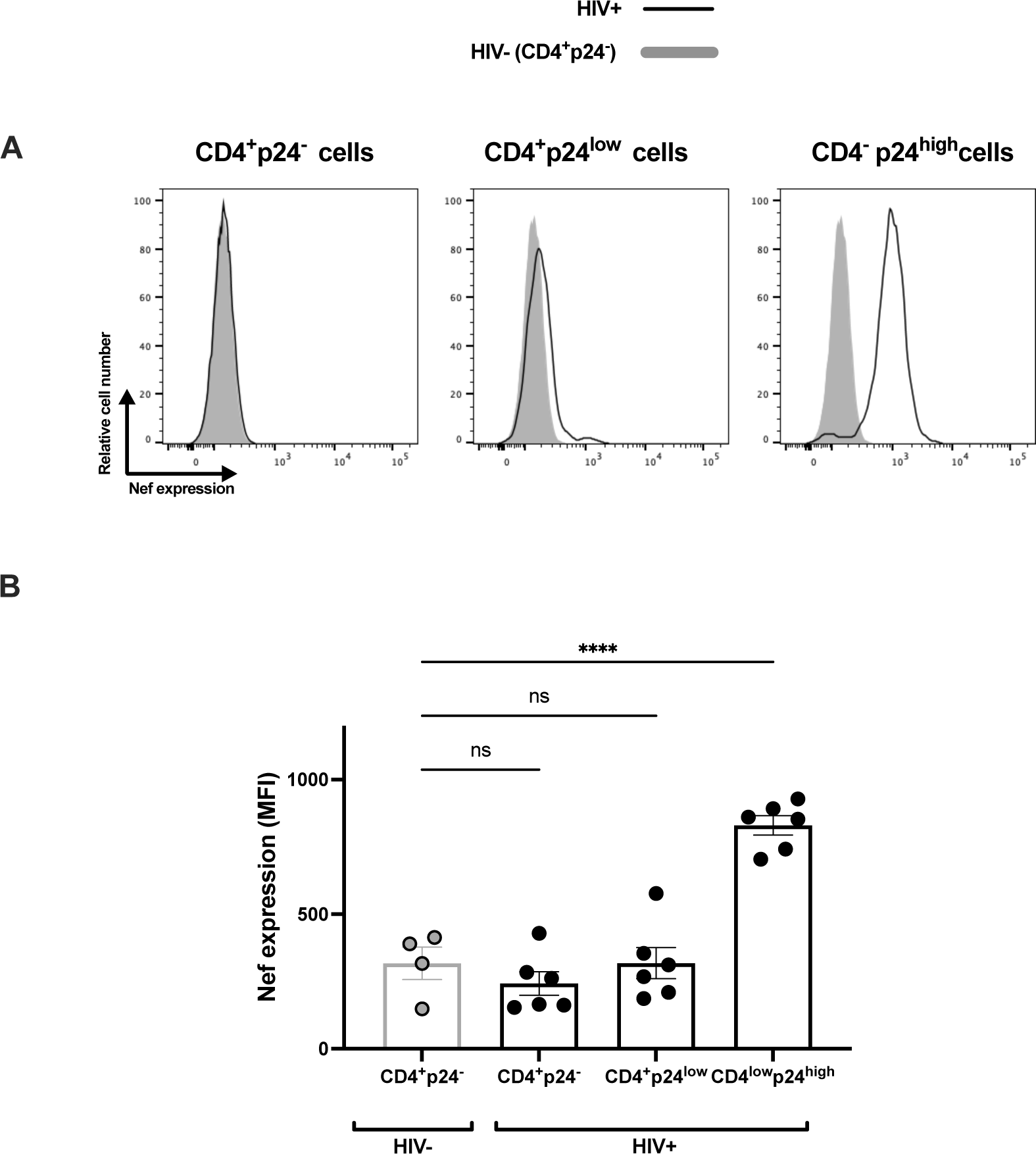
Nef expression in *ex vivo* expanded CD4+ T cells isolated from PLWH. *Ex vivo* expanded CD4 T cells from 6 HIV+ individuals and 4 HIV-individuals were stained for surface CD4 prior to detection of intracellular Nef and p24. (A) Histograms depicting representative staining. (B) Median fluorescence intensities (MFI) obtained with 6 HIV+ individuals and 4 HIV-individuals. Statistical significance was tested using one-way ANOVA test with a Holm-Sidak post-test (**** p<0.0001, ns: non-significant).

**Figure S6.**
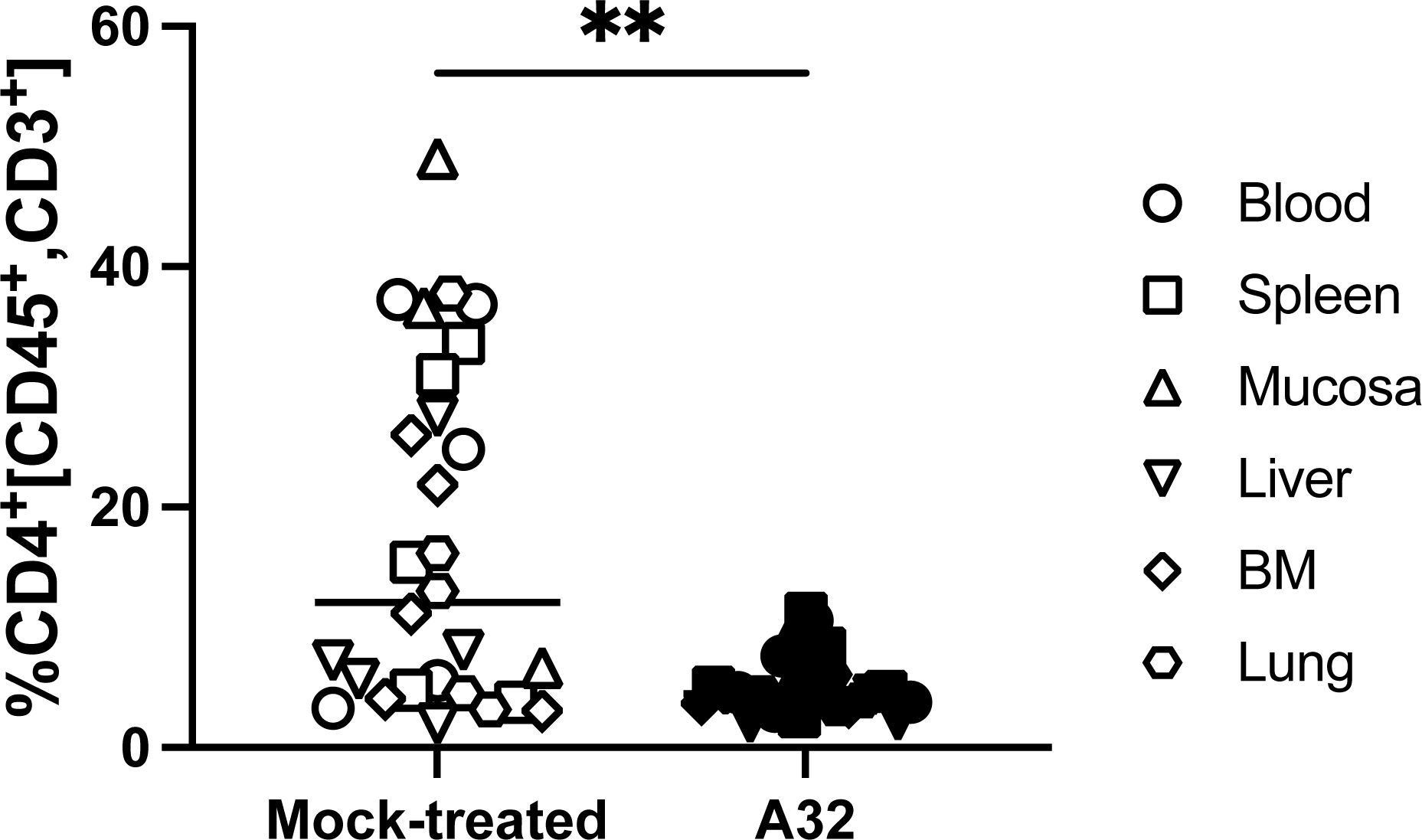
A32 reduces the levels of CD4+ T cells in tissues *in vivo*. NSG-15-Hu-PBL mice were infected with HIV-1 JRCSF intraperitoneally. At day 6 and 9 post infection, mice were administered 1.5 mg of A32 mAb subcutaneously (s.c.). The percentage of CD4+ T cells was evaluated by flow cytometry in blood, spleen, mucosa, liver, bone marrow (BM) and lung at day 11 post-infection. Statistical significance was tested using a Mann Whitney U test (** p<0.01).

